# Dynamics of the formation of flat clathrin lattices in response to growth factor stimulus

**DOI:** 10.1101/2025.05.22.655576

**Authors:** Lingxia Qiao, Marco A. Alfonzo-Méndez, Justin W. Taraska, Padmini Rangamani

## Abstract

Clathrin assemblies on the cell membrane are critical for endocytosis and signal transduction in cells. Specifically, Ω-shaped clathrin assemblies function as the coat of endocytic vesicles, while flat clathrin assemblies, also known as flat clathrin lattices, serve as signaling hubs for various signaling pathways. Multiple flat clathrin lattices exist on the cell membrane, and these lattices grow after epidermal growth factor stimulation (EGF) and then return to baseline. In this work, we used a particle-based model to simulate the assembly and disassembly of flat clathrin lattices to capture these dynamics. We found that the formation of flat clathrin lattices is highly dynamic, that is, cluster number, size and dwelling time often change even in the absence of any stimulus. Moreover, these key features are affected by adaptor protein 2 (AP-2) number, clathrin-clathrin binding rate, and clathrin diffusion coefficient. Specifically, an increase in AP-2 number leads to the transition from no cluster, short-lasting multiple small clusters, to a long-lasting single giant cluster. An increased clathrin-clathrin binding rate or decreased clathrin diffusion coefficient both result in an increased cluster number, reduced cluster size, and shortened dwelling time. Furthermore, we also predicted that under EGF stimulation, simultaneous changes in the AP-2 number, the clathrin-clathrin binding rate, and the clathrin diffusion coefficient can reproduce the experimentally observed trend of FCLs: an increase in cluster number and size in the first 30 minutes, followed by a decrease after 30 minutes. These findings reveal kinetic mechanisms underlying the formation of multiple FCLs and how EGF regulates FCL dynamics.

## 1 Introduction

Clathrin, a protein shaped like a triskelion, composed of three clathrin heavy chains and three light chains, plays an important role in growth factor signaling, adhesion, and endocytosis [1, 2]. Clathrin assembles into organized clusters on the cell membrane. One widely observed shape of clathrin assembles is the Ω-shape on the plasma membrane (PM) during endocytosis [3]. During endocytosis, clathrin molecules are recruited to the plasma membrane (PM) by adaptor protein complex 2 (AP-2), a major clathrin-associated adaptor [4]. Subsequently, the associated membrane bends into Ω-shaped pits [2]. In addition to the classic Ω shape pits, clathrin can also form a flat lattice structure on the PM (also termed plaques) [5]. Unlike Ω-shaped clathrin assemblies, flat clathrin lattices (FCLs) are long-lived stable structures [6, 7]. Recent studies have revealed that FCLs play an important role in cell functions. FCLs are consistently associated with actin [8] and have been shown to work with actin to oppose cell migration and contribute to skeletal muscle sarcomere organization [9, 10]. In addition, FCLs are enriched with β5-integrin [11, 12, 10], a receptor that plays a key role in cell proliferation and cell adhesion [13, 14]. Moreover, FCLs also regulate cell signaling by interacting with several signaling pathways, such as the epidermal growth factor (EGF), AKT, and hepatocyte growth factor pathways [9, 15, 16, 17].

Many mathematical models have been widely used to explore the mechanism of Ω-shaped clathrin in the absence or presence of cell membranes. In the absence of cell membranes, clathrins can still form a cage shape (the closed geometry of Ω shape) [18], and the clathrin is usually modeled as a coarse-grained triskelion particle whose interactions are controlled by the potential energy [19, 20, 21, 22]. However, clathrins closely interact with the cell membrane through adaptor proteins to form the Ω-shaped pits during endocytosis, thus inspiring the models that incorporate the cell membrane to the Ω-shaped clathrin [4, 1, 23, 24, 25]. For systems focusing on the biochemical aspects of the cell membrane, such as lipid localization and adaptor recruitment, clathrin is usually modeled as a coarse-grained triskelion particle, and computational approaches include Brownian dynamics with potential-based interactions [26, 27, 18, 28] and single-particle reaction-diffusion models [29, 30, 31]. For systems focusing on the biophysical aspects of cell membrane (e.g., tension and bending rigidity), clathrin is simplified to a zero-volume dot, and continuum field-based mesoscale models are preferred [25, 32, 33, 24, 34]. The involvement of the cell membrane opens the possibility to answer broader physiologically relevant questions such as the role of adaptors, cytoskeleton, and volume-to-area ratio in Ω-shaped clathrin, and the effect of clathrin dynamics on endocytosis.

Most of the models for Ω-shaped clathrin can be modified to models for the formation of FCL including particle-based molecular dynamics or Brownian dynamics with potential-based interactions [35], single-particle reaction-diffusion models [36, 30]. By changing the clathrin rotation [36] or strain energy [30], the simulated shape of clathrin can vary between Ω and lattice structure. Another way to acquire the clathrin lattice is to constrain the simulation domain to 2D [37, 35]. Furthermore, the 2D simulation domain can be discretized to a hexagonal lattice, and one clathrin occupies six edges of two neighboring hexagons, i.e., the lattice model [37]. Through simulations of these models, the formation of FCL has been reproduced, and the role of kinetics, for example, adaptor stoichiometry, clathrin concentration, and volume-to-area ratio, have been demonstrated [35, 37, 36, 30].

The computational studies mentioned above primarily focused on a single FCL. However, multiple FCLs exist on the cell membrane (Figure 1A). The distance between the two nearest FCLs can vary, as shown by the various distances between two FCLs in Figure 1B. The mean size of FCL in HeLa and U87 cells is smaller than 0.1μm^2^ (Figure 1C). Furthermore, for each FCL, there usually exists another FCL in the neighboring 1μm × 1μm square (Figure 1D). Compared with HeLa and U87 cells, MCF7 cells exhibit a larger number and size of FCLs on the cell membrane (Figure S1). Thus, FLCs are ubiquitous, having been observed in multiple cells from different origins [38, 17]. However, it remains unclear how neighboring multiple FCLs can co-exist instead of merging into one big cluster.

**Figure 1:**
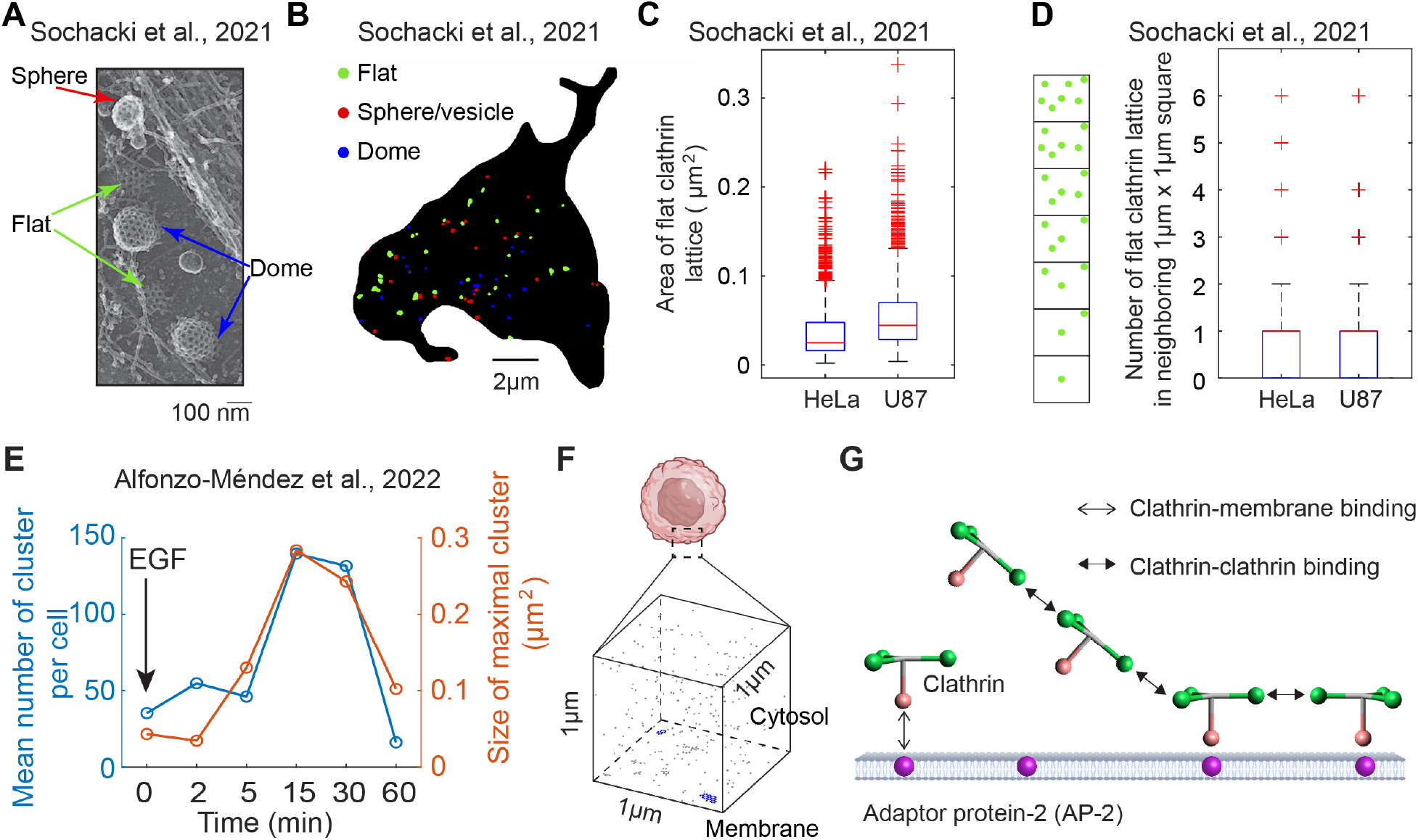
Multiple flat clathrin lattices (FCLs) coexist and undergo dynamic change under EGF stimulus. (A) A representative crop from a the montaged platinum replica transmission electron microscopy (PREM) image of an unroofed HeLa cell. This panel was adopted from Figure 1D in [38] [To editor: the permission is pending]. (B) Clathrin masks based on the PREM image of an unroofed HeLa cell. Clathrin clusters in flat, spherical, and dome shapes are indicated by green, red, and blue, respectively. Scale bar: 2*μ*m. Data are from [38]. (C) Area of flat clathrin lattices for HeLa cells (N=18) and U87 cells (N=12). In each box plot, the central red line denotes the median; bottom and top black edges indicate the 25th and 75th percentiles, respectively; whiskers denote the minimum and maximum extreme data points; plus markers represent outliers. Data are from [38]. (D) Number of FCLs in neighboring 1 *μ*m × 1*μ*m square. We used each FCL as a center and hen calculated the number of FCLs in the neighboring 1*μ*m × 1*μ*m square. The box plots were plotted in the same way as in (C). The data is based on the same HeLa cells (N=18) and U87 cells (N=12) as those in (C). (E) The total number of all FCLs (left axis) and size of maximal FCL (right axis) as a function of time under EGF stimulus. Figures are based on the data in [16]. (F) The cubic simulation domain. The bottom surface denotes membrane, and the area above denotes the cytosol. (G) Particle-based model that describes the binding and unbinding events of adaptor protein 2 (AP-2) and clathrins. This model is similar to that in [30]. The AP-2 (purple sphere) is located on the cell membrane. Clathrin (particle composed of one pink head and three green legs) can bind to AP-2 or other clathrins.

Moreover, multiple FCLs exhibit dynamic changes under EGF stimulus. Under the stimulus of the epidermal growth factor (EGF), FCLs on cellular membranes have been observed to grow in size and number, followed by a decrease toward baseline by 60 min (Figure 1E and [16]). More precisely, in the first 15 minutes after the stimulus of EGF, the number of FCLs increases nearly 3.5-fold (from 40 to 140), and maximal FCL area on the cell membrane grows near 6-fold (from 0.05μm^2^ to 0.3μm^2^). It should be noted that the increase in the size and number of FCLs occurs simultaneously. During this process, EGFR, scaffold protein Grb2, and tyrosine kinase Src are all recruited to FCLs ([16]). However, the underlying mechanism of such FCL dynamic changes remains unclear.

The above discoveries of multiple FCLs and the dynamics of EGF-triggered FCLs raise interesting questions about the biochemical mechanisms that govern the interaction between clathrins and the cell membrane. For example, how does clathrin form multiple stable clusters? What changes to kinetic parameters are required after EGF stimulation to achieve the observed behavior, that is, an initial increase in both the number and size of clusters in the first 30 minutes, followed by a return to baseline levels? Here, we sought to answer these questions using computational modeling.

## 2 Model development

In order to model the structure of flat clathrin lattice explicitly, we first chose a cube with edge length 1*μm* (one-twentieth of the diameter of HeLa cell [39]) as the simulation domain (Figure 1F). The bottom surface denotes a patch of the cell membrane and the volume above denotes the cytosol close to this membrane patch. Thus, we are able to simulate the exchange of clathrins between cell membrane and cytosol.

Since AP-2 is involved in the formation of FCLs by recruiting clathrin from the cytosol to the cell membrane, we included AP-2 in our simulation. Thus, we considered the diffusion and biochemical interactions of both clathrin and AP-2. AP-2 and clathrin are assumed to be located on the cell membrane and in the cytosol, respectively. However, clathrin can bind to the cell membrane by binding to AP-2 through the AP-2 binding site on clathrin. Moreover, other clathrins can bind to the AP-2 bound clathrin once these two clathrins are close enough, which allows the emergence of clathrin clusters. All of these association events are reversible. In addition to biochemical reactions, clathrins and AP-2 can diffuse, including translational and rotational diffusion. Clathrin can diffuse in the cytosol or on the cell membrane, while AP-2 can diffuse only on the cell membrane. Kinetic parameters and diffusion coefficients are from [30] (also see Table S1). In [30], kinetic parameters were optimized to fit the fold change of clathrin cluster size in *in vitro* experiments [40], and diffusion coefficients were estimated from the Stokes-Einstein equation.

Next, we describe how clathrin and AP-2 are modeled. Each clathrin is modeled by a rigid structure (in green and pink in Figure 1G) that can diffuse in the cytosol or on the membrane. For each clathrin, there are four binding sites (see Table S2 for the length of binding sites): three binding sites (green in Figure 1G) that allow it to bind to another clathrin, and one binding site (pink in Figure 1G) that allows clathrin to bind to AP-2. This structure of clathrin is the same as that in [30] except that the binding site for AP-2 is decreased to 1. This simplification is used to reduce computation time, and we assume that it does not quantitatively affect the results. Moreover, the clathrin-clathrin binding event is assumed to be achieved by head to head binding of clathrin binding sites from two different clathrins. After binding, the distance between two bound sites is 5 nm, and the center of two clathrins and all clathrin binding sites are on the same plane. For the binding of clathrin to AP-2, the AP-2 binding site on the clathrin is oriented perpendicular to the cell membrane. The excluded volume of clathrin is set at 10 nm. These settings ensure the formation of a flat hexagonal structure of clathrins on the cell membrane.−

Then, to model the dynamics of clathrins in the cubic simulation domain, we utilized the particle-based algorithm NERDSS, a nonequilibrium simulator for multibody self-assembly developed by Varga et al. [31]. The time step Δ*t* is set to 3μs, during which we assumed that each clathrin either diffuses or undergoes only one reaction. Within each time step, the dissociation events are tested first. The corresponding reaction probability of the dissociation event is calculated as 1 − *exp*(−*k*Δ*t*) from a Poisson process, where *k* is the dissociation rate with the unit s^−1^. Then, the association events are tested, where the reaction probability is calculated by free propagator reweighting (FPR) algorithm [36, 41, 31]. If the particle undergoes dissociation or association events, this particle cannot diffuse in this time step. Finally, the particles that are not involved in any reactions in this time step will diffuse. The assembled clathrin cluster will diffuse as an entire unit with the following diffusion coefficient: 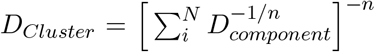. Here, *N* is the total number of components; *n* is 1 for the translational diffusion coefficient and 3 for the rotational diffusion coefficient. During the diffusion process, the displacement of the particle is calculated as 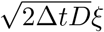, where *D* is the diffusion coefficient and *ξ* is a random number generated from a standard normal distribution, respectively. At the same time, the algorithm checks whether the newly displaced positions will cause overlapping of the clathrin molecules. If overlap occurs, the displacement is resampled until non-overlapping positions among the clathrins are achieved.

The input files used for NERDSS are available at https://github.com/RangamaniLabUCSD/FlatClathrinLattice. The simulations were performed at Triton Shared Computing Cluster at the San Diego Supercomputer Center (https://doi.org/10.57873/T34W2R).

## 3 Results

In this study, we investigated the underlying mechanisms of two phenomena: (1) the formation of multiple FCLs in the absence of stimulation, and (2) the EGF-induced dynamics of FCLs, characterized by an increase in both their size and number, followed by a return to baseline levels. We utilized NERDSS, a nonequilibrium reaction-diffusion self-assembly simulator [30], to describe the assembly and disassembly of clathrins. In this model, each clathrin is an individual particle and interacts with each other through binding and unbinding events (Figure 1F-G). We found that even if all the kinetic parameters remain unchanged, FCLs are very dynamic, meaning the number and size of FCLs often vary with time. The numbers of FCLs, size of each FCL, and dwelling time are affected by adaptor protein-2 (AP-2) number, clathrin-clathrin binding rate, and clathrin diffusion coefficient. Furthermore, simultaneous changes in AP-2 number, clathrin-clathrin binding rate, and clathrin diffusion coefficient can generate the experimentally observed FCLs dynamics triggered by EGF, that is, the increasing trend of cluster number and size in the first 30 minutes and then decreasing back to baseline levels. These findings reveal the formation mechanisms of multiple FCLs and provide insights into the mechanisms underlying EGF-triggered FCL dynamics, enhancing our understanding of how FCLs form and change on the cell membrane.

### 3.1 The formation of flat clathrin lattices is dynamic

Given the presence of multiple FCLs in various cell types even without stimulus (Figure 1A-D and [38]), we first studied the mechanism of spontaneous formation multiple FCLs, corresponding to the case without any stimulus. We set the randomly distributed clathrins as the initial condition (Figure 2A), and then simulated clathrin dynamics. We found that the cluster is always dynamic: there is one cluster at 21 minutes but two clusters at 29.7 minutes (Figure 2B). The mean cluster size, defined as the total cluster size divided by the cluster number, also changes from 0.075 *μm*^2^ to 0.0285 *μm*^2^. Such dynamic transitions in cluster number (blue curve in Figure 2C) occur frequently and persists even when the number of AP-2-bound clathrins reaches the plateau (red curve in Figure 2C). These results indicate that FCLs on the cell membrane can transition between different states.

**Figure 2:**
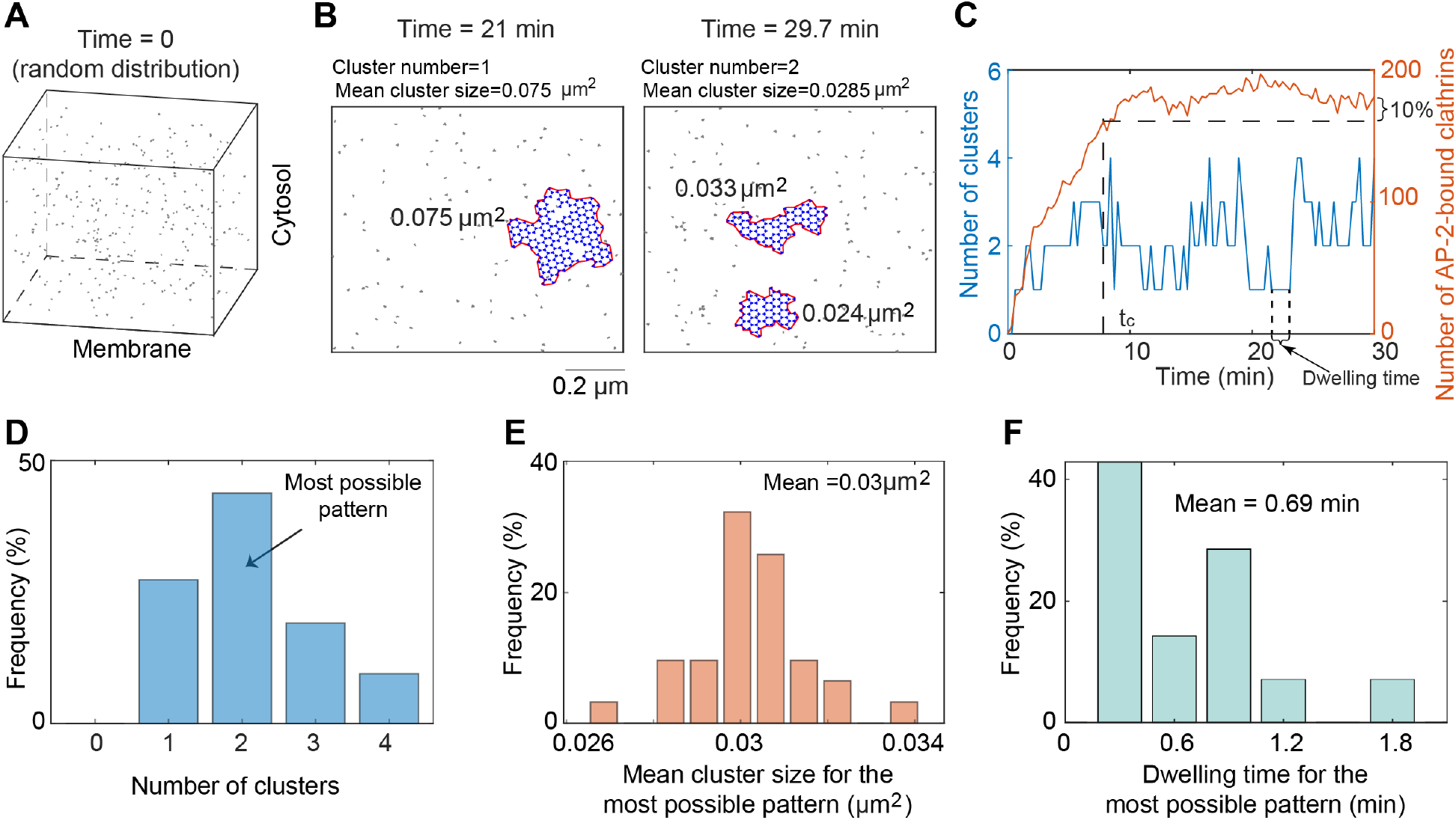
The FCL formation is highly dynamic. (A) The initial condition. Clathrin molecules are randomly distributed in the simulation domain at time 0. (B) Snapshots of FCLs at 21 minutes (left) and 29.7 minutes (right) when simulating from the random initial condition in (A). The number of AP-2 is set to be 150. The area surrounded by the red curve is calculated as the size of each cluster. Mean cluster size is defined as the total size of all clusters divided by the number of clusters at a specific time point. (C) The number of clusters (left axis) and that of AP-2 bound clathrins on cell membrane (right axis) as functions of time. The settings of model simulations are the same as that in (B). We defined *t*_*c*_ as the time when the number of clathrins on the cell membrane first exceeds 90% of the amount present at 30 minutes, and assumed that the system after *t*_*c*_ is at equilibrium. (D) The frequency of cluster numbers after *t*_*c*_. The cluster number with the highest frequency is defined as the most possible pattern. (E) The frequency of mean cluster size for the most possible pattern. (F) The frequency of dwelling time for the most possible pattern.

To quantitatively measure the FCL dynamics on the cell membrane, we defined three metrics for the most possible pattern: cluster number, mean value of the mean cluster size, and mean value of dwell time. First, we defined *t*_*c*_ as the time when the number of AP-2-bound clathrins reaches 90% of that at the end of simulations (i.e., 30 minutes). We chose the time 30 minutes due to the long-lived property of FCL which ranges from 2 to 10 min to more than 1 h [6, 11]. We only focused on the behavior after *t*_*c*_ (Figure 2C), because the number of AP-2-bound clathrins only shows small fluctuations after *t*_*c*_. Next, we calculated the frequency for different cluster numbers after *t*_*c*_, and define the pattern with the highest frequency of cluster number as the most possible pattern (Figure 2D). For example, for the simulation in Figure 2B, the most possible pattern has a cluster number of 2. However, the system at different time points may show same cluster number, but the mean cluster size is not the same. Thus, we focused on the most possible pattern, and calculated the mean value of the mean cluster size (Figure 2E). The mean value reflects the size of individual cluster that appears frequently. Similarly, we calculated the mean of the dwelling time for the most possible pattern (Figure 2F) Here, the dwelling time is the duration that the cluster number remains unchanged (arrow in Figure 2C). For example, for the simulation in Figure 2B, the most possible pattern has a mean cluster size of 0.03μm^2^ and a dwelling time of 0.69 minutes. Overall, these results suggest that the most possible pattern exhibits frequent changes in cluster number, size, and dwelling time in the absence of any stimulus.

### 3.2 Number of AP-2 affects the cluster number, size and dwelling time

Previous studies showed that sufficient adaptor proteins are required to maintain the FCL [30], but the effect of AP-2 number on the cluster number, size, and dwell time remains unknown. To investigate this, we simulated the clathrin dynamics for different numbers of AP-2. The initial condition is the same as that in Figure 2A, that is, randomly distributed clathrins. Despite the dynamic transition between different states, the number of clusters is 0, larger than 1, and 1 when AP-2 number is 10, 200, and 400, respectively (Figure 3A-C). By testing more values of the AP-2 number with 4 replicates for each value, the cluster number for the most possible pattern shows a significant increase and then decrease when the AP-2 number increases (Figure 3D). These results indicate that increasing AP-2 number leads to the transition from no cluster, multiple small clusters to one giant cluster. It should be noted that we used a higher clathrin-clathrin binding rate than that in [30]. When the clathrin-clathrin binding rate is low, the clathrin behavior with increased AP-2 number is the same as that in [30], that is, a transition directly from no cluster to one giant cluster without experiencing the multiple small clusters stage (Figure S2). In addition to the effect on cluster number, the AP-2 number also affects the mean cluster size and dwelling time. As the number of AP-2 increases, the mean value of the mean cluster size monotonically increases (Figure 3E). At the same time, the dwelling time for the most possible pattern first decreases and then increases (Figure 3F), which is opposite to the trend of the cluster number. Combining the effects of AP-2 number on cluster number, cluster size, and dwell time, we concluded that increasing the number of AP-2 leads to the transition from no cluster to short-lived multiple small clusters, to a long-lasting single giant cluster.

**Figure 3:**
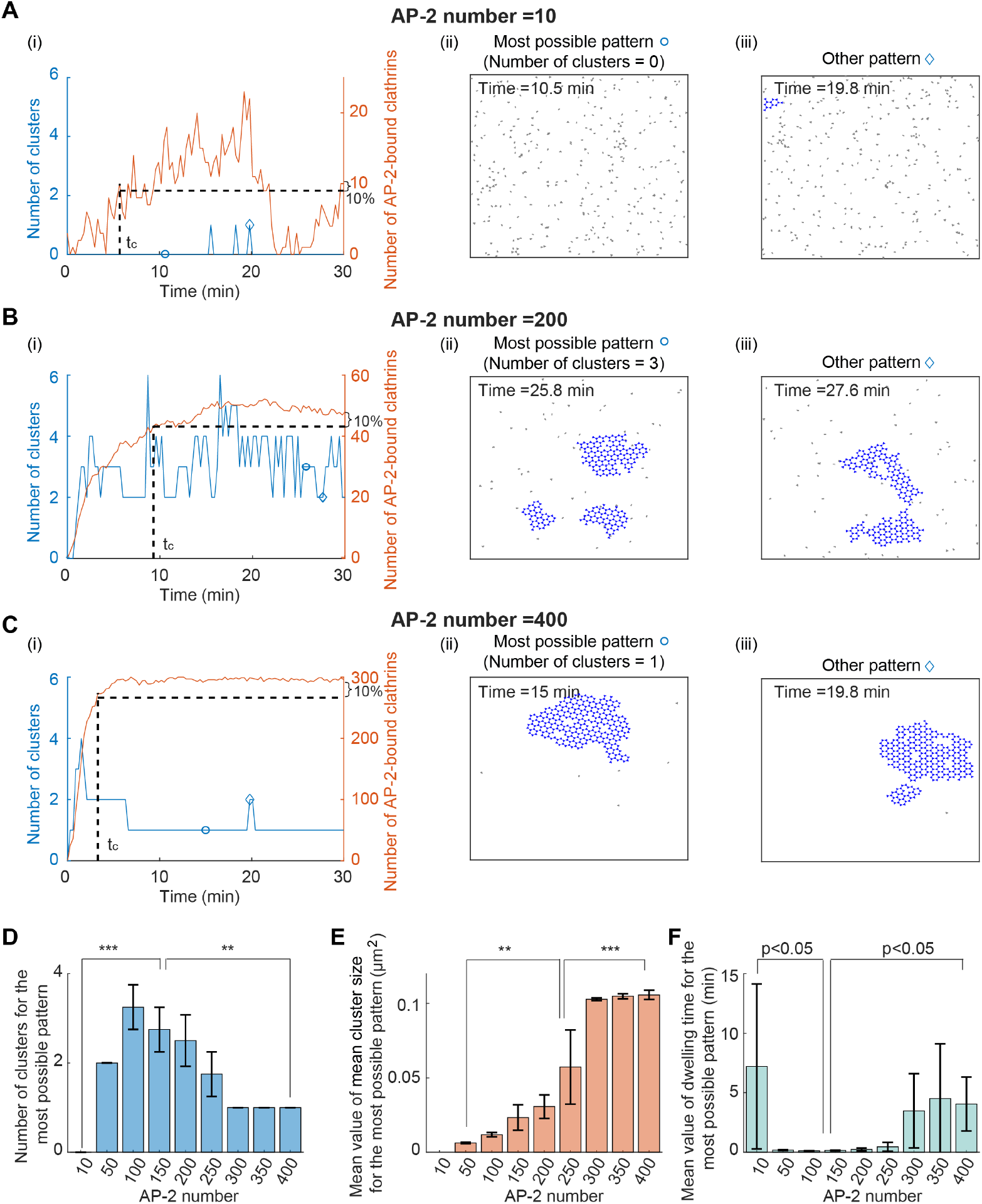
Number of AP-2 affects the FCLs cluster number, size and dwelling time. (A-C) Simulated clathrin dynamics when the AP-2 number is 10 (A), 200 (B), and 400 (C). The initial condition is randomly distributed clathrins in the simulation domain. In panel (i), the number of clusters (in blue) and the number of AP-2 bound clathrins (in red) are shown as a function of time. Panels (ii) and (iii) show the most possible pattern and the other pattern, respectively. The circle or diamond indicates the time point in the panel (i). (D) The cluster number for the most possible pattern when increasing the AP-2 number. (E) The mean value of the mean cluster size for the most possible pattern when increasing the AP-2 number. (F) The mean dwelling time for the most possible pattern when increasing the AP-2 number. P value was obtained from unpaired t-test: ** indicates p<0.01;*** indicates p<0.001. Data in (D-F) were shown as mean ± SD, where SD means standard deviation.

### 3.3. Increasing clathrin-clathrin binding rate leads to an increased cluster number, reduced cluster size, and shortened dwelling time

Next, we studied the influence of clathrin-clathrin binding rate on cluster formation. We changed the value of clathrin-clathrin binding rate *k*_(*AP* −2*·*)*Clat·Clat*_ while fixing the value of other kinetic parameters. The initial condition is still a random distribution of clathrins. As the clathrin-clathrin binding rate *k*_(*AP* −2*·*)*Clat·Clat*_ increases, the most possible pattern shows increased cluster number (Figure 4A-D), decreased mean value of the mean cluster size (Figure 4E), and decreased mean dwelling time (Figure 4F). This result suggests that a high clathrin-clathrin binding rate leads to an increased cluster number, reduced cluster size, and shortened dwelling time.

**Figure 4:**
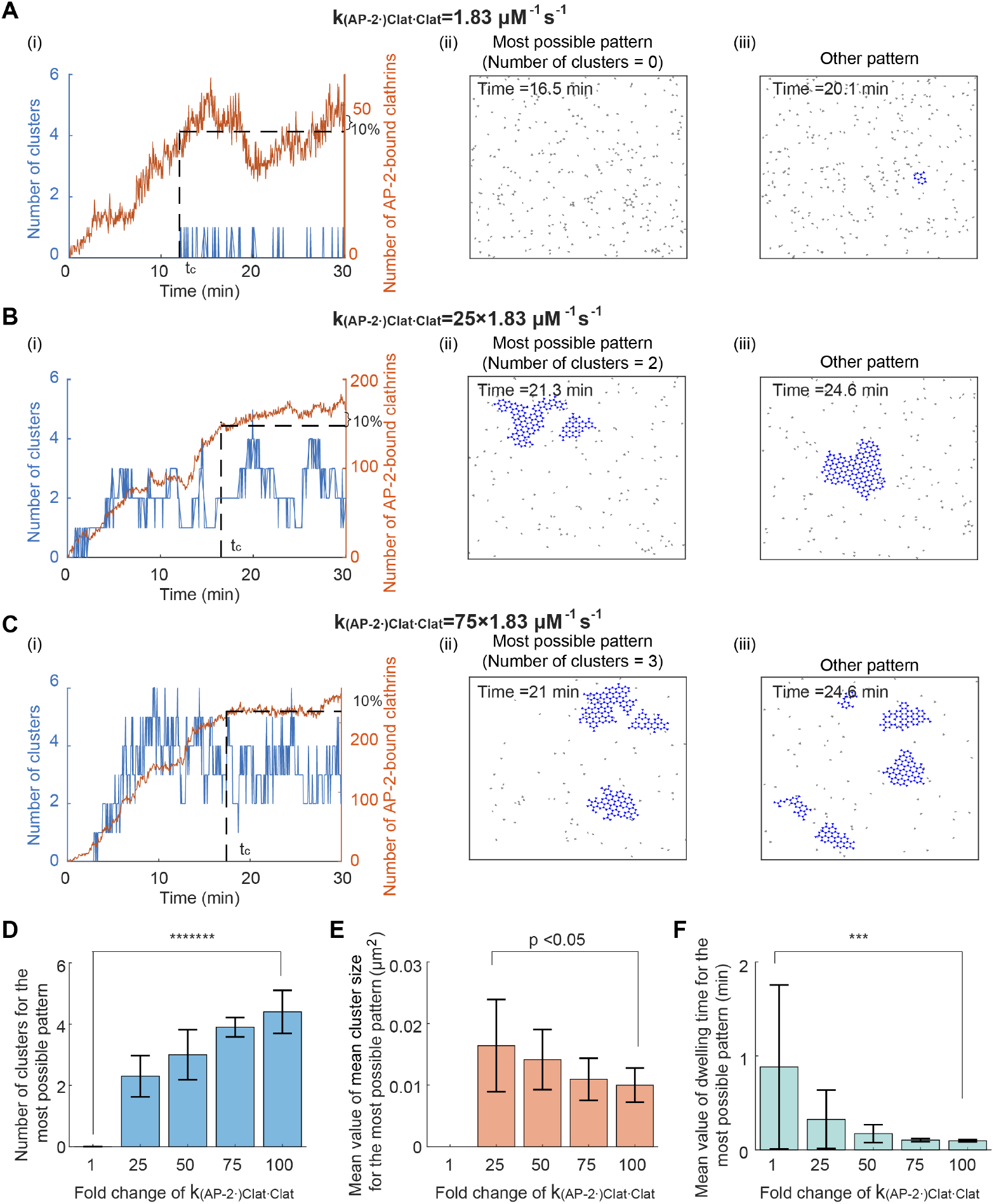
Increasing clathrin-clathrin binding rate leads to an increased cluster number, reduced cluster size, and shortened dwelling time. (A-C) Simulated clathrin dynamics when the clathrin-clathrin binding rate *k*_(*AP* −2*·*)*Clat·Clat*_ is 1.83*μ*M^−1^s^−1^ (A), 25×1.83*μ*M^−1^s^−1^ (B), and 75×1.83*μ*M^−1^s^−1^ (C). Parameters except *k*_(*AP* −2*·*)*Clat·Clat*_ are fixed. The initial condition is that clathrins are randomly distributed in the simulation domain. Panels (i-iii) are plotted in the same way as those in Figure 3A. (D) The cluster number for the most possible pattern when increasing *k*_(*AP* −2*·*)*Clat·Clat*_. (E) The mean value of the mean cluster size for the most possible pattern when increasing *k*_(*AP* −2*·*)*Clat·Clat*_. (F) The mean dwelling time for the most possible pattern when increasing *k*_(*AP* −2*·*)*Clat·Clat*_.P value was obtained from unpaired t-test: *** indicates p<0.001;******* indicates p<1E-7. Data in (D-F) were shown as mean ± SD, where SD means standard deviation.

### 3.4 Decreasing clathrin diffusion coefficient results in an increased cluster number, reduced cluster size, and shortened dwelling time

Next, we investigated how the diffusion coefficient of clathrin affects the formation of multiple clusters. Similar to how we identified the role of clathrin-clathrin binding rate, we changed the clathrin diffusion coefficient but fixed other kinetic parameters. We found that, when the clathrin diffusion coefficient decreases, the most possible pattern exhibits a similar trend to those observed with increased *k*_(*AP* −2*·*)*Clat·Clat*_: increased cluster number, decreased cluster size, and decreased dwelling time (Figure 5). Thus, a low clathrin diffusion coefficient results in an increased cluster number, reduced cluster size, and shortened dwelling time.

**Figure 5:**
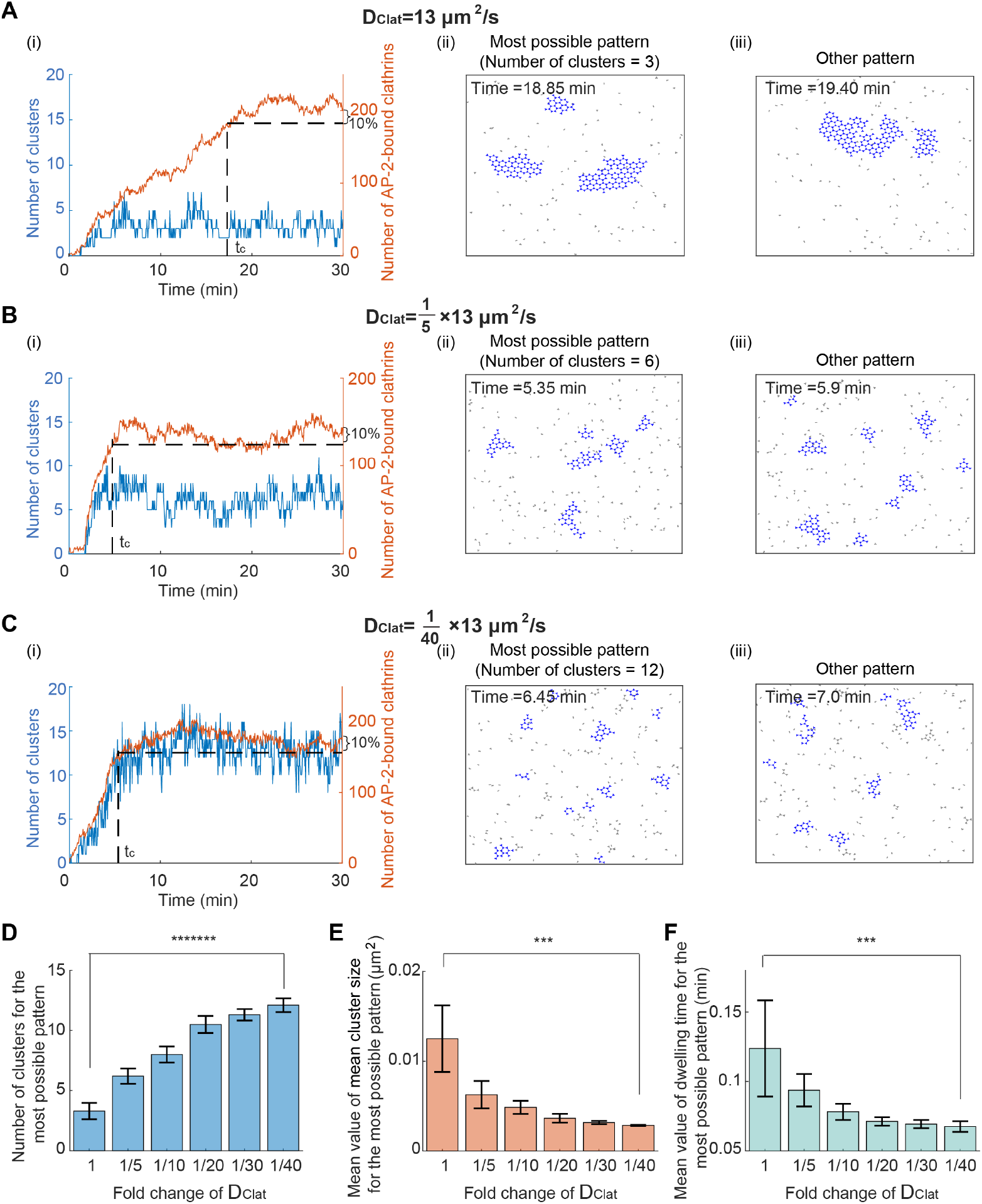
Decreasing clathrin diffusion coefficient results in an increased cluster number, reduced cluster size, and shortened dwelling time. (A-C) Simulated clathrin dynamics when the clathrin diffusion coefficient *D*_*Clat*_ is 13*μ*m^2^/s [30] (A), 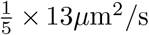 (B), and 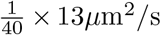 (C). Parameters except *D*_*Clat*_ are fixed. The initial condition is that clathrins are randomly distributed in the simulation domain. Panels (i-iii) are plotted in the same way as those in Figure 3A. (D) The cluster number for the most possible pattern when decreasing *D*_*Clat*_. (E) The mean value of the mean cluster size for the most possible pattern when decreasing *D*_*Clat*_. (F) The mean dwelling time for the most possible pattern when decreasing *D*_*Clat*_. P value was obtained from unpaired t-test: *** indicates p*<*0.001;******* indicates p*<*1E-7. Data in (D-F) were shown as mean*±*SD, where SD means standard deviation.

### 3.5 Simultaneous changes in the clathrin-clathrin binding rate and clathrin diffusion coefficient can reproduce the experimentally observed trend of FCLs under EGF stimulus

The above studies focus on clathrin behavior without stimulus, that is, all kinetic parameters in any given simulation are fixed. In this case, although both the clathrin number and size are dynamic after a long time simulation (e.g., after *t*_*c*_), their mean behavior remains relatively stable, deviating from the experimentally observed increasing trend during the 15 minutes after the EGF stimulus. Therefore, we hypothesized that kinetic parameters associated with the assembly of FCL might change in response to an EGF stimulus. To test this hypothesis, we started from a state where there is only one cluster (Figure 6A), and then tested different combinations of parameter changes (Figure 6B) to determine which one best reproduces the experimental results.

**Figure 6:**
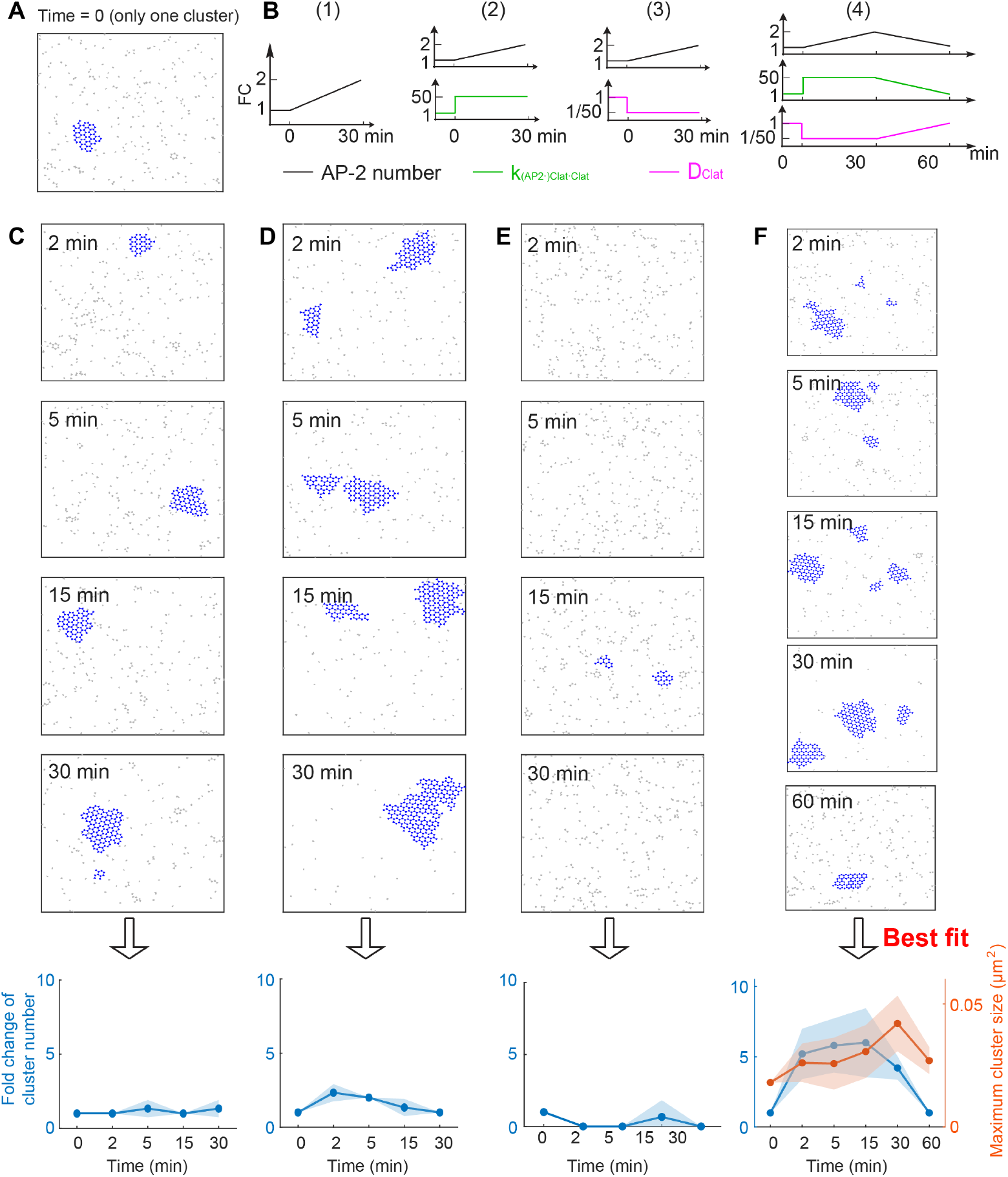
Simultaneous changes in the clathrin-clathrin binding rate and clathrin diffusion coefficient can reproduce the experimentally observed trend of FCLs under EGF stimulus. (A) Initial condition used to mimic the clathrin cluster before EGF stimulus. (B) Changing kinetic parameters to mimic the effect of EGF stimulus. Four different combinations were tested: (1) increasing AP-2 number only; (2) increasing AP-2 number and clathrin-clathrin binding rate *k*_(*AP* −2*·*)*Clat·Clat*_ simultaneously; (3) increasing AP-2 number and decreasing clathrin diffusion coefficient *D*_*Clat*_ simultaneously; (4) increasing AP-2 number, increasing *k*_(*AP* −2*·*)*Clat·Clat*_, and decreasing *D*_*Clat*_ simultaneously. (C-F) Clathrin dynamics on the cell membrane under different parameter changes in (B). The panels (C-F) correspond to parameter changes in (1-4) in the panel (B), respectively. For each type of parameter change, four or five snapshots of clathrins at different time points were shown. The last column shows the fold change in cluster number (left) and maximum cluster size (right) as a function of time. For each type of parameter change, three replicates were performed, shown as mean *±* SD (standard deviation). Solid lines in the last row of (C-F) denote the mean, with the shading representing the standard deviation.

First, we varied the kinetic parameters to explore which configurations could reproduce the experimentally observed clathrin dynamics during 30 minutes after the EGF stimulus. In this time interval, experimental data showed that both the cluster number and cluster size increase (Figure 1E and [16]). As shown in the previous sections, increasing AP-2 number alone, increasing clathrin-clathrin binding rate *k*_(*AP* −2*·*)*Clat·Clat*_ alone, or decreasing clathrin diffusion coefficient *D*_*Clat*_ alone all lead to the increased cluster number. Thus, in order to generate the increased cluster number during 30 minutes after EGF stimulus, these three parameters might be changed, but it remains unknown whether these three parameters should be changed individually or simultaneously. Thus, we tested the following four ways of changing parameter (Figure 6B): (1) increasing the AP-2 number; (2) increasing the AP-2 number and increasing clathrin-clathrin binding rate *k*_(*AP* −2*·*)*Clat·Clat*_ simultaneously; (3) increasing the AP-2 number and decreasing clathrin diffusion coefficient *D*_*Clat*_ simultaneously; (4) increasing AP-2 number, increasing *k*_(*AP* −2*·*)*Clat·Clat*_, and decreasing *D*_*Clat*_ simultaneously. For all methods, AP-2 number is increased from 100 to 200 in the time interval [0, 30 minutes]; *k*_(*AP* −2*·*)*Clat·Clat*_ is increased by a 50-fold change; *D*_*Clat*_ is decreased to one-fiftieth of its previous value. We found that the clathrin dynamics in the first three methods do not show a significant increase in cluster number in the time interval [0, 30 minutes] (Figure 6C-E). However, the fourth method, where AP-2 number, *k*_(*AP* −2*·*)*Clat·Clat*_ and *D*_*Clat*_ are all changed, exhibits an increased cluster number and increased cluster size in the time interval [0, 30 minutes] (Figure 6F). These results indicate that a simultaneous increase in AP-2 number and *k*_(*AP* −2*·*)*Clat·Clat*_, along with a decrease in *D*_*Clat*_, can generate the experimentally observed cluster dynamics during the first 30 minutes of the EGF stimulus.

After we reproduced the clathrin dynamics in the first 30 minutes of EGF stimulus, we hypothesized that reversing all the kinetics parameters would reproduce the experimentally observed clathrin disappearance between 30-60 min after EGF stimulation. We observed that decreasing AP-2 number, *k*_(*AP* −2*·*)*Clat·Clat*_ and *D*_*Clat*_ lead to decreases in both FCLs cluster number and size at 60 min of EGF stimulation (Figure 6F). This result is consistent with the experimental data collected after 60 minutes of EGF stimulation (Figure 1E). Between 30 and 60 minutes, numerical simulations predicted that large clusters break apart into many smaller clusters (Figure S3) and thus result in a temporary increase in the number of clusters. These results also suggest that reverting the AP-2 number, *k*_(*AP* −2*·*)*Clat·Clat*_, and *D*_*Clat*_ to their baseline values reproduces the clathrin dynamics during 30 to 60 minutes after EGF stimulus.

## 4. Discussion

Clathrin can assemble into various shapes, including Ω-shaped pits and flat clathrin lattices. The Ω-shaped pits that are formed on the cell membrane during endocytosis are short-lived and are removed from the membrane after the process. In contrast, flat clathrin lattices tend to be more stable, lasting more than 10 minutes, and have different functions compared to the Ω-shaped pits. These flat clathrin lattices have been found to act as signaling hubs. For example,their number and size exhibit an increasing trend followed by a decreasing trend after the EGF stimulus. However, how such changes in number and size are achieved remains elusive. Here, we investigated the mechanisms underlying the formation of multiple FCLs, as well as the changes in FCLs induced by EGF stimulus.

We used a particle-based model for clathrins and simulated the dynamics using NERDSS, a nonequilibrium simulator for multibody self-assembly [31]. For the mechanism of multiple FCLs formation, we started from a random distribution of clathrins and only focused on the clathrin state when the system becomes “stable”, that is, the time when the number of AP-2 bound clathrins reaches 90% of that at the end of the simulation. We revealed that the FCL formation is dynamic, suggested by the highly dynamic cluster number and cluster size. Moreover, when increasing AP-2 number, clathrin states exhibit a transition from no cluster to short-lasting multiple small clusters, and finally to long-lasting single giant cluster. An increase in the clathrin-clathrin binding rate or a decrease in the clathrin diffusion coefficient both lead to an increased cluster number, a decreased cluster size, and reduced dwelling time. In addition to the mechanism of multiple FCL formation, we also studied the mechanisms of changes in FCLs caused by EGF stimulus. We found that the changes in AP-2 number are not sufficient to achieve the experimentally observed trend of FCLs. However, when combined with changes in clathrin-clathrin binding rate and clathrin diffusion coefficient, the simulated clathrin dynamics qualitatively fit the experimental data.

The roles of kinetic parameters in multiple FCLs formation found in this work improve our understanding of clathrin assemblies. We found that increasing clathrin-clathrin binding rate leads to an increased cluster number, reduced cluster size, and shortened dwelling time. Such an increase in the number of clusters may be because a high value of *k*_(*AP* −2*·Clat·Clat*)_ allows formed clusters to easily absorb unbound clathrins when the component occasionally dissociates. Furthermore, as more clusters arise, they compete for the limited clathrin resources, potentially resulting in a decrease in cluster size. As small clusters do not have too many clathrins and may lose all their clathrins in a very short time, small clusters tend to disappear quickly, which may cause a decrease in dwelling time. As for the role of clathrin diffusion coefficient, we showed that decreasing clathrin diffusion coefficient results in an increased cluster number, reduced cluster size, and shortened dwelling time. This can be explained by the fact that a slow clathrin diffusion coefficient restricts the dissociated clathrin from dispersing too far, thus allowing the clathrin clusters to recruit the dissociated clathrin to maintain their components effectively instead of merging into a giant cluster or disassembling. This also could be related to the relatively new hypothesis of “frustrated endocytosis” that proposes that long-lived FCLs arise from adhesive forces generated from integrins that physically prevent clathrin from curving [42]. However, further experiments are required to validate these simulation results.

The model predictions are consistent with previous biochemical evidence to some extent. On the one hand, we predicted that the increase in the AP-2 number helps FCLs grow after EGF stimulus. This is closely related to the experimentally observed increased interaction between clathrin and endocytic adaptors (e.g., AP-2, Eps15) after EGF stimulus [43, 44, 45]. Moreover, there is a regulatory domain on the EGFR cytoplasmic domain that is essential for AP-2 recruitment [46]. On the other hand, we predicted that the change in clathrin-clathrin binding rate might be important for EGF-triggered FCLs dynamics. Such changes may relate to EGF-triggered phosphorylation of AP-2 and clathrin [47, 48]. Further experimental work is necessary to validate this hypothesis.

FCLs regulate various cellular processes like proliferation and actin network remodeling [9, 49]. FCLs also harbor different receptors including epidermal growth factor receptor (EGFR), fibroblast growth factor receptor 1 (FGFR1), low-density lipopoprotein receptor LDLR, lysophosphatidic acid receptor 1 (LPAR1), and β5 integrin [16, 50]. β5-integrin knock down leads to decreased abundance of FCLs at the plasma membrane [16]. Therefore, flat clathrin lattices serve as multifunctional platforms that coordinate signaling, endocytosis, and cytoskeletal organization. Despite the distinct dynamic behaviors of FCLs under various stimuli, their assembly and disassembly may universally be highly dynamic.

In addition to the particle-based model we used in this work, lattice model [51, 52], Cahn–Hilliard equation [53, 54, 55] Turing model [56, 57], molecular dynamics simulations [58, 59] have also been used to model the protein clusters within the cell. However, the continuous models, including Cahn–Hilliard equation and Turing model, do not include the stochasticity of chemical reactions, and thus may lead to the failure of capturing experimental observations. This failure was also validated by simulating Turing model (Figure S4), where the increases in cluster number and size occur at different times. Thus, when simulating the dynamics of multiple clusters, continuous models may not be able to fit experiments as well as particle-based models.

In this work, we focused on the interactions between AP-2 and clathrins with several simplifications. For example, there are multiple AP-2 binding sites on clathrin [60], but we assume only one AP-2 binding site per cluster for the sake of simplicity; the binding between AP-2 and the cell membrane is neglected by assuming that AP-2 is located on the cell membrane at all times. These simplifications are not expected to qualitatively affect the results, as the model still captures the phenomenon of AP-2-clathrin binding at the plasma membrane. Another limitation is that we did not consider the reactions between clathrins and signaling molecules involved in EGFR pathway. In this study, we predicted that, to ensure the increasing trend of FCL number and size in the first 30 minutes and then decreasing trend between 30 and 60 minutes, EGFR pathway will regulate the AP-2 number, clathrin-clathrin binding rate, and clathrin diffusion coefficient simultaneously. However, the mechanism of how EGFR pathway regulates these parameters remains unclear. Future work is expected to include both EGFR pathways and clathrins to investigate the crosstalk between the EGFR pathway and FCLs. Furthermore, there are debates about the relationship between flat clathrin lattices and Ω-shaped pits [2, 61, 11, 62], which were not considered in this work. Both constant curvature and constant area models have been used to explain the transition from flat to curved clathrin assemblies [11]. To further investigate the crosstalk among flat clathrin lattices, Ω-shaped pits, and EGFR pathways, which still remain elusive, future modeling studies will need to develop a model that incorporates membrane properties, self-assembled clathrins, and EGFR signaling pathways.

The protein assemblies studied in this work have been widely observed within the cell. On the cell membrane, there exist many clusters formed by receptors or scaffold proteins, for example, nanodomain formed by A-kinase anchoring protein (AKAP) 79/150 [63], clusters formed by AMPA- and NMDA-type glutamate receptors[64]. In the cytosol, several intrinsically disordered proteins could also form assemblies (also known as biomolecular condensates or liquid-like droplets), including stress granule [65] and condensates formed by type I regulatory subunit of cAMP-dependent protein kinase (RIα) [66]. In the nucleus, there are also condensates that control gene regulations, for instance, Yes-associated protein (YAP) [67] condensates and PML nuclear bodies [68]. Researchers have extensively explored the formation mechanisms of these protein assemblies [69, 70]. For example, increased monomer concentration leads to the transition from co-existing multiple assemblies to one giant assembly [51], consistent with our simulation results when increasing AP-2 number. xt); the lag time and steepness for the clathrin cluster growth curve are primarily controlled by the individual clathrin binding rate to membrane and the binding rate between clathrin and AP-2, respectively [30]; two-dimensions mesoscopic simulations allow the formation of multiple postsynaptic protein assemblies while three-dimensions cannot [71]. In addition to formation mechanisms, the interaction between protein assemblies and neighboring environment also has been widely studied [70, 72, 73]. For instance, protein assemblies not only regulate key signaling pathways [74, 75] but also affect the kinetic parameters [76]; curvature-inducing proteins can assemble and regulate cell membrane shape [77]. This study also serves as an example for investigating the interactions between protein assemblies and their surrounding environment.

## 5 Acknowledgment

P.R. was supported by NIH R01GM132106 and Army Research Office W911NF231249. J.W.T. was supported by the Intramural Research Program of the National Heart, Lung, and Blood Institute, National Institutes of Health, USA.

## Supplement

### 1 The Turing model cannot simultaneously achieve the increase of cluster size and the increase of cluster number

Motivated by the Turing model in [56], we used the following equations to describe the dynamics of AP-2 and clathrin on the cell membrane (Figure S4A):

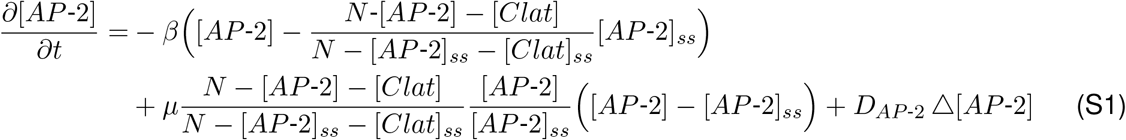

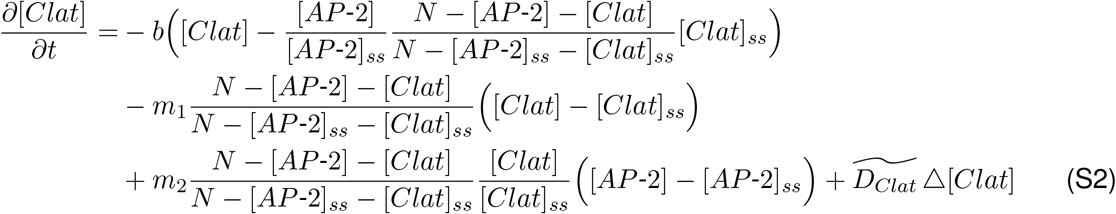

where [*AP* −2] and [*Clat*] denote the concentrations of AP-2 and clathrin on the cell membrane, respectively. All terms in Equations (S1) and (S2) are the same as those in [56] except the diffusion terms. The diffusion terms *D*_*AP* −2_ △[*AP* −2] and 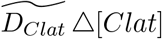 in Equations (S1) and (S2) are simpler than that in [56], but the Turing pattern still can be maintained. Four mechanisms that are key to Turing pattern formation are as follows: 1) the enhanced recruitment of AP-2 if one AP-2 has already bound to the cell membrane (the curved arrow in Figure S4A); 2) the recruitment of clathrin to the cell membrane caused by AP-2 (the arrow from AP-2 to clathrin in Figure S4A); 3) the steric repulsion between clathrin and AP-2 (the arrow from clathrin to AP-2 in Figure S4A); 4) much slower diffusion coefficient of AP-2 compared to that of clathrin. In the first two mechanisms, the reaction strength is denoted by *μ* and *m*_2_. For the third mechanism, it is achieved by multiplying 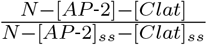 to each reaction term: the larger the [*AP* −2] + [*Clat*] is, the smaller the 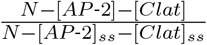 is, leading to less recruitment events. Here, *N* is set to be a large enough number to ensure [*AP* −2] + [*Clat*] ≤ *N* hold all the time, and the subscript *ss* denotes the homogeneous steady-state value. As for the fourth mechanism, we set the ratio of diffusion coefficients between AP-2 and clathrin 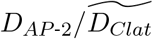 as 0.0377 in our simulations.

The values of kinetic parameters in Equations (S1) and (S2) are listed in Table S3, which are determined by searching previous clathrin models or the requirement of Turing instability. According to the clathrin model in [30], the dissociation rate of AP-2 and PIP_2_ is 1 s^−1^, and the dissociation rate of clathrin and AP-2 is 0.03 s^−1^. Therefore, we set the dissociation rate of AP-2 and cell membrane *β* to be 1 s^−1^, and the dissociation rate of clathrin and cell membrane *b* 0.03 s^−1^. Besides, the values of *N*, [*AP* −2]_*ss*_ and [*Clat*]_*ss*_ were chosen to match the scale of AP-2 in [30] (~ 361 copies/*μ*m^2^). Other kinetic parameters were determined by the constraints of Turing instability, such as *μ, m*_1_, *m*_2_, *D*_*AP* −2_ and 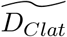. It should be noted that the value of *D*_*AP* −2_ is 1% of AP-2 translational diffusion constant on the membrane in [30], and that the value of 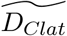 is also 1% of clathrin translational diffusion constant in [30]. This rescaling of the diffusion constant ensures the correct size of each clathrin cluster.

We first tested whether the Turing model can capture the experimental observations of FCL dynamics after the stimulus of EGF. By simulating the Turing model, we obtained a stable pattern, corresponding to the system without stimulus (the first plot in Figure S4B). Then, we increased the association rate of AP-2 and membrane *μ* by 80% to mimic the effect of adding EGF. Under this change, the system experiences two stages: (1) in [0, 15 seconds], each clathrin cluster grows in size but no new clusters emerge (Figure S4B); (2) after 15 seconds, each large clathrin cluster disassembles into small clusters, leading to the increase in the number of clusters and decrease in the size of each cluster. To make the trend of cluster size and cluster number more clear, we used the image processing tool to obtain the boundary of each cluster and then calculated the cluster size and cluster number quantitatively (Figure S4C). It can be seen that the increase in cluster size and the increase in total cluster number occur at different time intervals, i.e., [0 15 seconds] and after 15 seconds, respectively. However, as we can see from the experimental data (Figure 1E), the increase in the size and that in the number happens simultaneously. Taken together, the Turing model cannot capture the FLC dynamics under the EGF stimulus.

**Figure S1:**
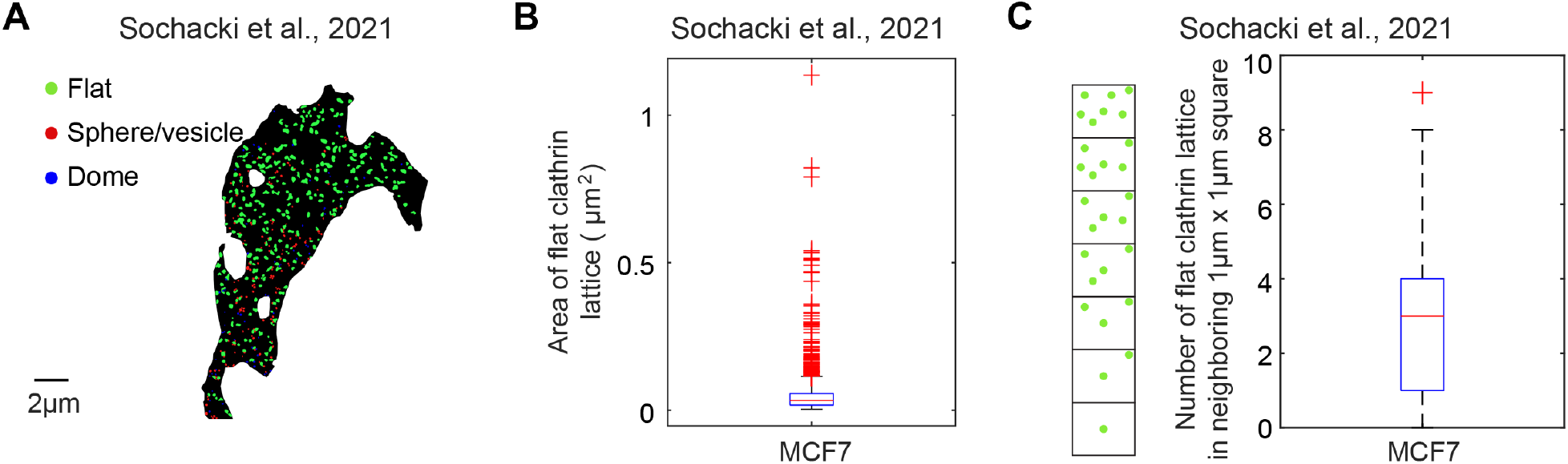
The FCL distributions in MCF7 cells. (A-C) The same plots as those in Figure 1B-D except the cell type.

**Figure S2:**
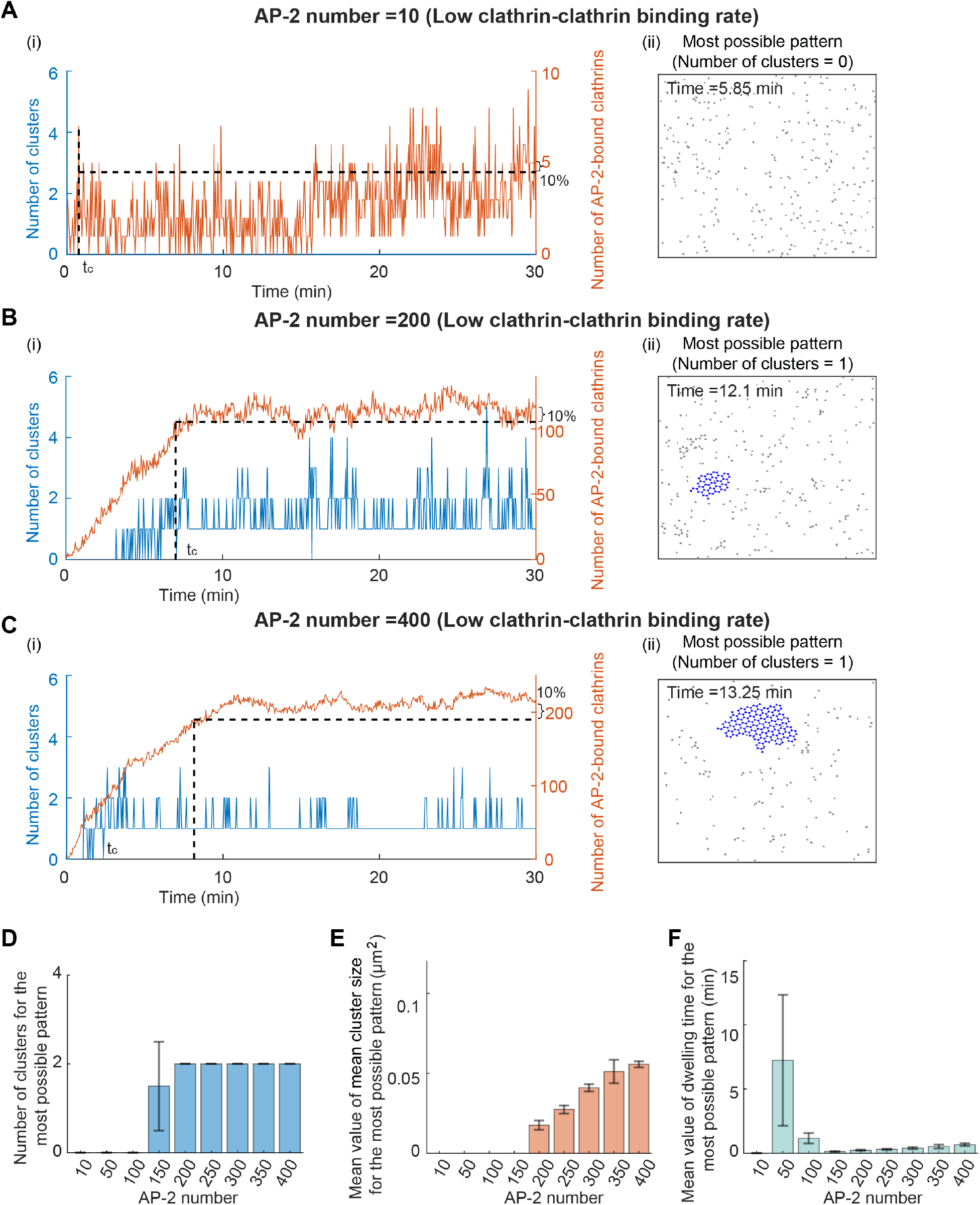
The FCL exhibits the phase transition from no cluster to one giant cluster when the AP-2 number increases while maintaining a low clathrin-clathrin binding rate. (A-F) The same plots as those in Figure 3 except that a low clathrin-clathrin binding rate is used.

**Figure S3:**
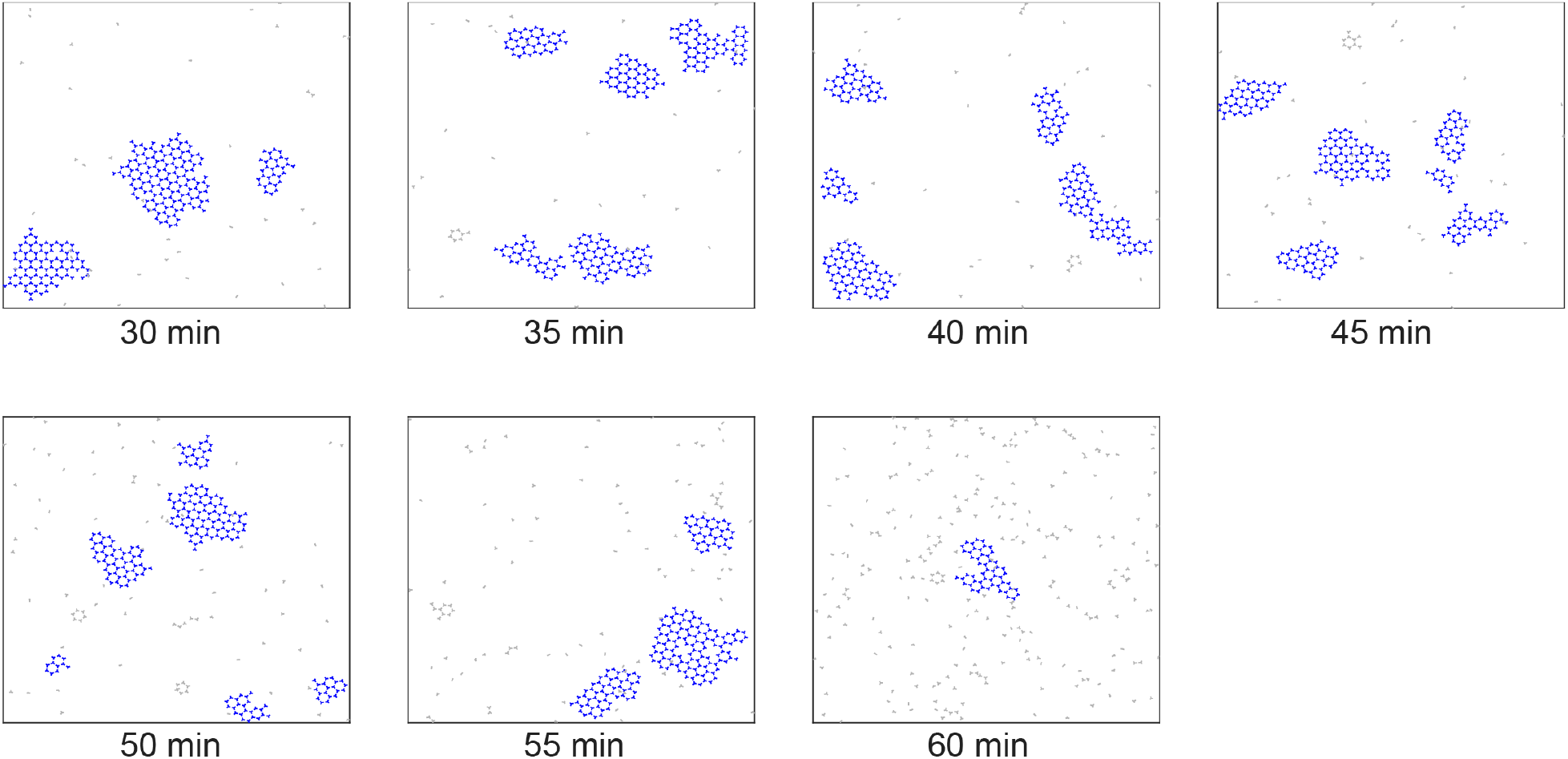
Simulated FCL dynamics after 30 minutes of EGF stimulus when kinetic parameters change follow the way in Figure 6B(4).

**Figure S4:**
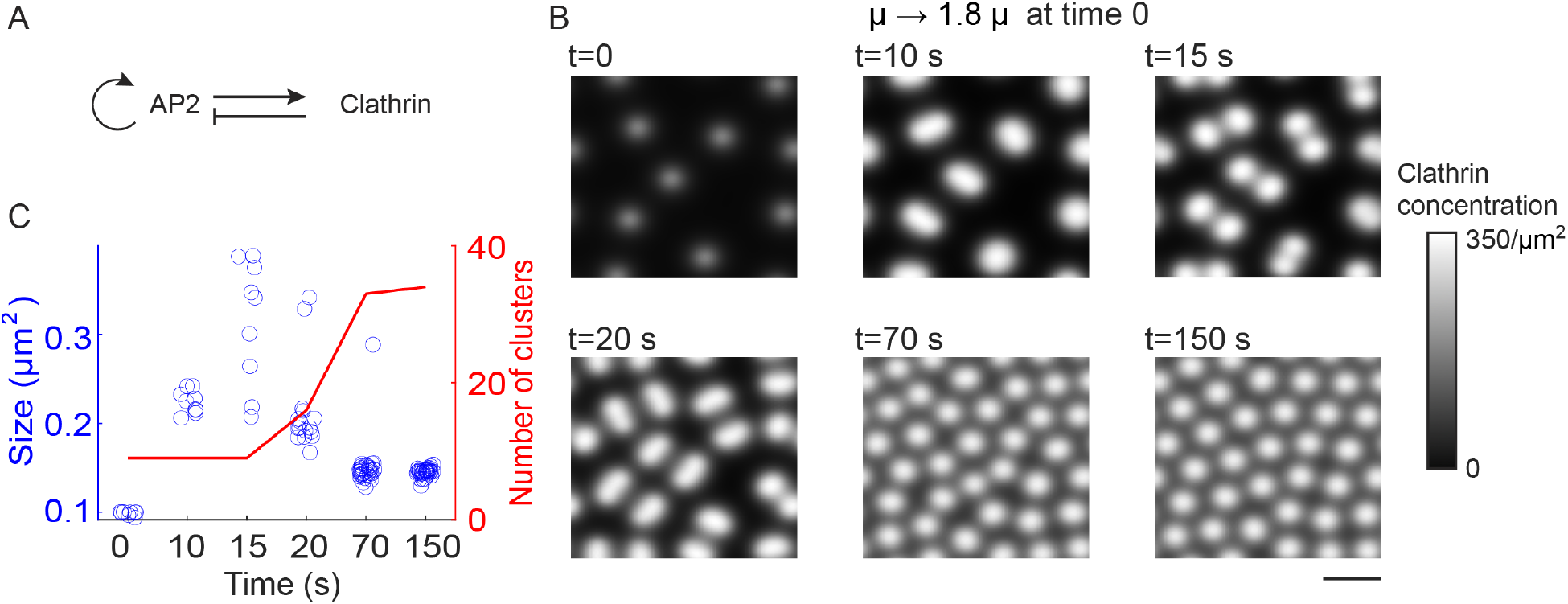
The Turing model cannot simultaneously achieve the increase of cluster size and the increase of cluster number. (A). Schematic of reactions between the AP-2 and the clathrin. The AP-2 improves the recruitment of itself and the clathrin to the cell membrane. Once bound to the cell membrane, the clathrin inhibits the accumulation of the AP-2 on the cell membrane due to steric repulsion. (B). The snapshots of clathrin clusters after increasing the association rate of AP-2 and membrane *μ* by 80% at time 0. The first plot corresponds to the Turing pattern with parameters in Table S3. Scale bar is 1 *μ*m. (C). The size for each clathrin cluster (left axis) and the total number of clathrin clusters (right axis) at the time points in (B). The increase in the clathrin cluster size only occurs between 0 and 15 seconds, while the increase of the total number of clathrin clusters occurs after 15 seconds.

**Table S1:**
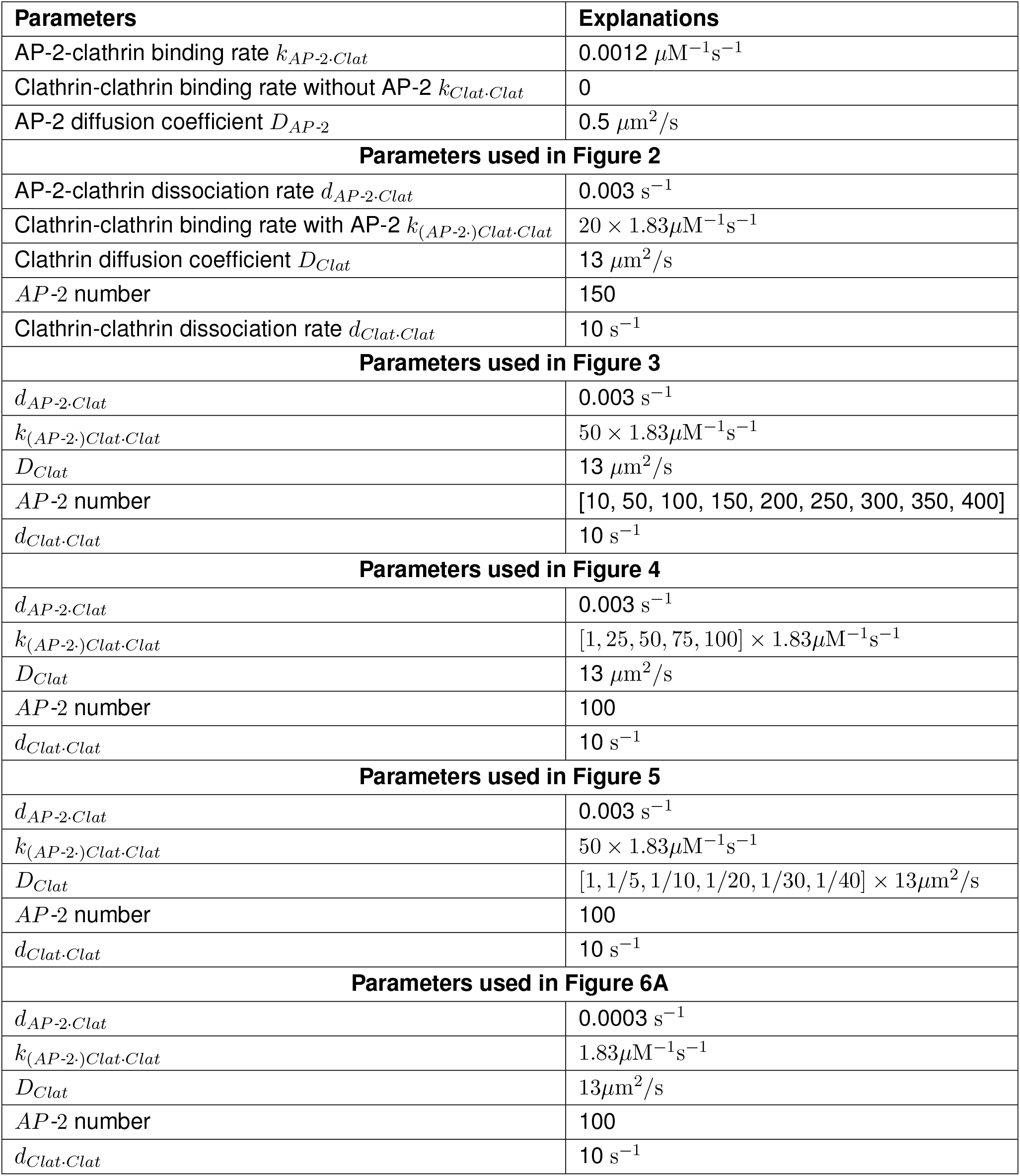
Parameters used in the particle-based model (from [30])

**Table S2:**
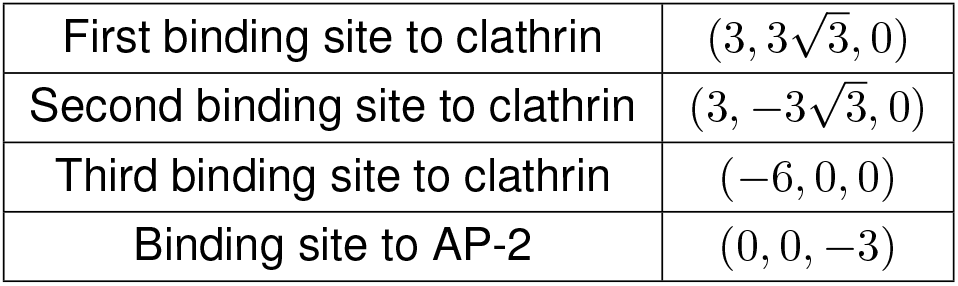
Relative coordinate for clathrin.

**Table S3:** Parameters used in Turing model.

| Parameters | Explanations | Values | Ref. |
| --- | --- | --- | --- |
| **Parameters used in the Turing model** |  |  |  |
| $\beta$ | AP-2 & membrane dissociation rate | $1 \text{ s}^{-1}$ | [30] |
| $\mu$ | AP-2 & membrane association rate | $1.4 \text{ s}^{-1}$ | Fitted |
| $b$ | Clathrin & membrane dissociation rate | $0.03 \text{ s}^{-1}$ | [30] |
| $m_1$ | Clathrin & membrane dissociation rate caused by unspecified molecules | $0.8 \text{ s}^{-1}$ | Fitted |
| $m_2$ | Recruitment rate of clathrin by AP-2 | $10 \text{ s}^{-1}$ | Fitted |
| $D_{AP-2}$ | Diffusion coefficient of AP-2 | $0.049 \mu\text{m}^2\text{s}^{-1}$ | 1/100 of the value in [30] |
| $\widetilde{D_{Clat}}$ | Diffusion coefficient of clathrin | $1.3 \mu\text{m}^2\text{s}^{-1}$ | 1/100 of the value in [30] |
| $N$ | Maximal total concentration | $500 \mu\text{m}^{-2}$ | Same scale as in [30] |
| $[AP-2]_{ss}$ | Homogeneous steady-state value of [AP-2] | $25 \mu\text{m}^{-2}$ | Same scale as in [30] |
| $[Clat]_{ss}$ | Homogeneous steady-state value of [Clat] | $25 \mu\text{m}^{-2}$ | Same scale as in [30] |

